# Compendium-wide analysis of *P. aeruginosa* core and accessory genes reveal more nuanced transcriptional patterns

**DOI:** 10.1101/2022.04.14.488429

**Authors:** Alexandra J. Lee, Georgia Doing, Samuel L. Neff, Taylor Reiter, Deborah A. Hogan, Casey S. Greene

## Abstract

Strains of *Pseudomonas aeruginosa*, an opportunistic pathogen that causes difficult to treat infections, have significant genomic heterogeneity including the presence of diverse accessory genes that are only present in some strains or clades. Both core genes, which are conserved across strains, and accessory genes have been associated with traits such as biofilm formation and virulence. Much of what we know about core and accessory gene content comes from genome analyses. Here, we use a newly assembled transcriptome compendium to analyze the transcriptional patterns of core and accessory gene expression in PAO1 and PA14 strains across thousands of samples from hundreds of distinct experiments. We found that a subset of core genes were stable, having consistent correlated expression patterns across samples regardless of strain background, with a focus on strains PAO1 and PA14. These stable core genes had fewer co-expressed neighbors that were accessory genes.

**Significance:** *Pseudomonas aeruginosa* is a ubiquitous pathogen. There is a lot of diversity amongst *P. aeruginosa* strains, some which are clinically relevant. Understanding how these different strain-level traits manifest is important for identifying targets that regulate different traits of interest. With the availability of a PAO1-mapped and PA14-mapped RNA-seq compendium, which contain hundreds of strains, it is now possible to examine the effect of different strains on expression, which can mediate different traits. In this study we developed an approach to compare expression profiles across different *P. aeruginosa* gene groups – core and accessory genes. This approach revealed a subset of core genes with different transcriptional patterns across strains, which could contribute to trait differences.

## Introduction

*Pseudomonas aeruginosa* is a gram-negative bacterium that is able to thrive in a variety of different abiotic environments including soil and water, and live in association with plants and animals.^1^ *P. aeruginosa* is also an opportunistic human pathogen that is frequently implicated in hospital-acquired infections^2,3^ and is a particular concern for immunosuppressed and vulnerable individuals^4,5^.

Amongst the *P. aeruginosa* strains, there is much phenotypic diversity. Clinical and environmental strains exhibit varying capacity for biofilm formation^6,7^, levels of antibiotic susceptibility^8,9^, metabolic profiles^10^ and differences in virulence factor production^11^. Some strains can also perform environmental biotransformations, which are chemical modifications.^12,13^ This Strain-level phenotypic diversity is reflected in its genetic diversity; the *P. aeruginosa* genome contains both conserved core genes and strain-specific accessory genes.^14,15^ A phylogenetic analysis across 1,311 strains, using only core genes, divided *P. aeruginosa* strains into five major lineages^14^. Within these five lineages there were two predominant lineages that most strains belonged to – PAO1 and PA14.^16^ Compared to strain PAO1, strain PA14 was found to be more virulent in a number of model systems.^17,18^ Comparative genomic studies found that this difference in virulence has been attributed to the presence or expression of accessory genes^17,19–22^ and core genes^18,21,23^. There were also distinct surface-sensing circuits that contributed to differences in cell surface attachment in different strain backgrounds.^24–26^ These data highlight how the same factors can be deployed in different ways.

Strain-level differences between PAO1 and PA14 lineages have predominantly been studied based on genetic composition.^17–21,23–26^ However, how genetic differences translate into differences in the transcriptome and ultimately phenotypes is not well understood. The many transcription factors (TFs) and other transcriptional regulatory elements found in *P. aeruginosa* control the expression of many gene products that mediate various different traits^27,28^ such as virulence^29–31^ and these can vary across strains. For example, a study by Sana *et al*. found a set of core genes that were differentially expressed in PAO1 compared to PA14.^32^ Many of these genes were involved in quorum sensing, which is a cell-cell signaling communication system that regulates many virulence factors.^33^ For another example, within the type III secretion system (T3SS), which is a virulence determinant that allows *P. aeruginosa* to deliver toxic effector proteins to host cells. Strains PAO1 and PA14 both secrete effectors ExoT and ExoY, but they also differ in that PAO1 secretes ExoS and PA14 secretes ExoU.^34^ The production of these accessory virulence factors was found to be regulated by strain-specific response regulators^35^ and shared regulators, like ExsA^36^. The secretion of ExoU was found to increase lethality in mice more than ExoS.^37^ Overall, ExoS and ExoU effect different host pathways.^38^

In general, transcriptional regulatory networks are known to be versatile across *P. aeruginosa* strains^35^; this versatility was observed across strains within a single species in other microbes as well^39–41^. These network differences between strains were due, in part, to the accessory genome – there were some strain-specific TFs, target genes (i.e. *exoS* and *exoU)* and TF-target gene interactions.^35,39,40^ Overall, the above examples demonstrated that both core and accessory regulation affect strain-level traits, like virulence. While the existence of the core and accessory genome has been known, how different genomic backgrounds affect transcriptional profiles remains to be explored. Extending the findings from Trouillon *et al*., who found that there existed some conserved and some different regulatory interactions between PAO1 and PA14, we can examine the downstream transcriptional patterns across PAO1 and PA14, accounting for the different gene groups.

In this paper we examined the transcriptional patterns within and between *P. aeruginosa* core and accessory genes using compendia of expression data from the two most profiled strain types, PAO1 and PA14 (Doing *et al*., co-submitted manuscript^42^). While most studies focused on only a single condition when studying core and accessory transcription^20,43–46^, we demonstrated that transcriptional patterns for core and accessory gene can be elucidated by comparing expression across many experiments performed by different labs which complements studies that examine individual experiments alone.^47–51^ Leveraging newly formed *P. aeruginosa* RNA-seq compendia created by Doing *et al*., co-submitted manuscript,^42^ in which 2,333 RNA-seq samples were aligned to both PAO1 and PA14 cDNA reference genomes, we found that amongst core genes, there exists a subset that exhibit stable transcriptional patterns across strain type, while others differ substantially. The most stable core genes are less often co-expressed with accessory genes compared to the least stable core genes. By enabling a more nuanced understanding of transcriptional behavior across *P. aeruginosa* strains, the RNA-Seq compendium helps to further elucidate differences in how genes in *P. aeruginosa* are regulated and contribute to phenotypic diversification.

## Results

### PAO1 and PA14 compendia contain a diverse collection of experiments

We generated transcriptomics compendia using 2,333 recently downloaded, filtered and normalized public *P. aeruginosa* RNA-seq samples assembled by Doing *et al*., co-submitted manuscript.^42^ All samples in the dataset were separately aligned to both a PAO1 cDNA reference and a PA14 cDNA reference genome, regardless of strain annotation. For accessory genes that were specific to either PAO1 or PA14, provided by the BACTOME website^52^, we calculated the median expression for each sample (Figure 1A). The PAO1 specific-gene set values were calculated using the PAO1-mapped RNA-seq compendium and the PA14 specific gene set values were calculated using the PA14-mapped RNA-seq compendium. When all samples were plotted based on median expression of PAO1-and PA14-specific genes, it was clear that most samples had high expression of only one strain-specific set. Strain annotations in SRA entries were present for approximately 70% of samples, and these annotations strongly supported the use of accessory gene expression as a predictor of strain identity (Figure 1A). A threshold was applied to the median expression of each set of strain-specific accessory genes. This threshold was determined based on the distribution of accessory gene expression in samples annotated as either PAO1 or PA14 (insets in Figure 1B and C, respectively) to identify whether other samples were obtained from strains that are either PAO1-like, PA14-like, or distinct from both (e.g., clinical isolates). Using these cutoffs, a strain PAO1 sample compendium, with 890 samples (Figure 1B), and a strain PA14 sample compendium, with 505 samples (Figure 1C), were created.

**Figure 1:**
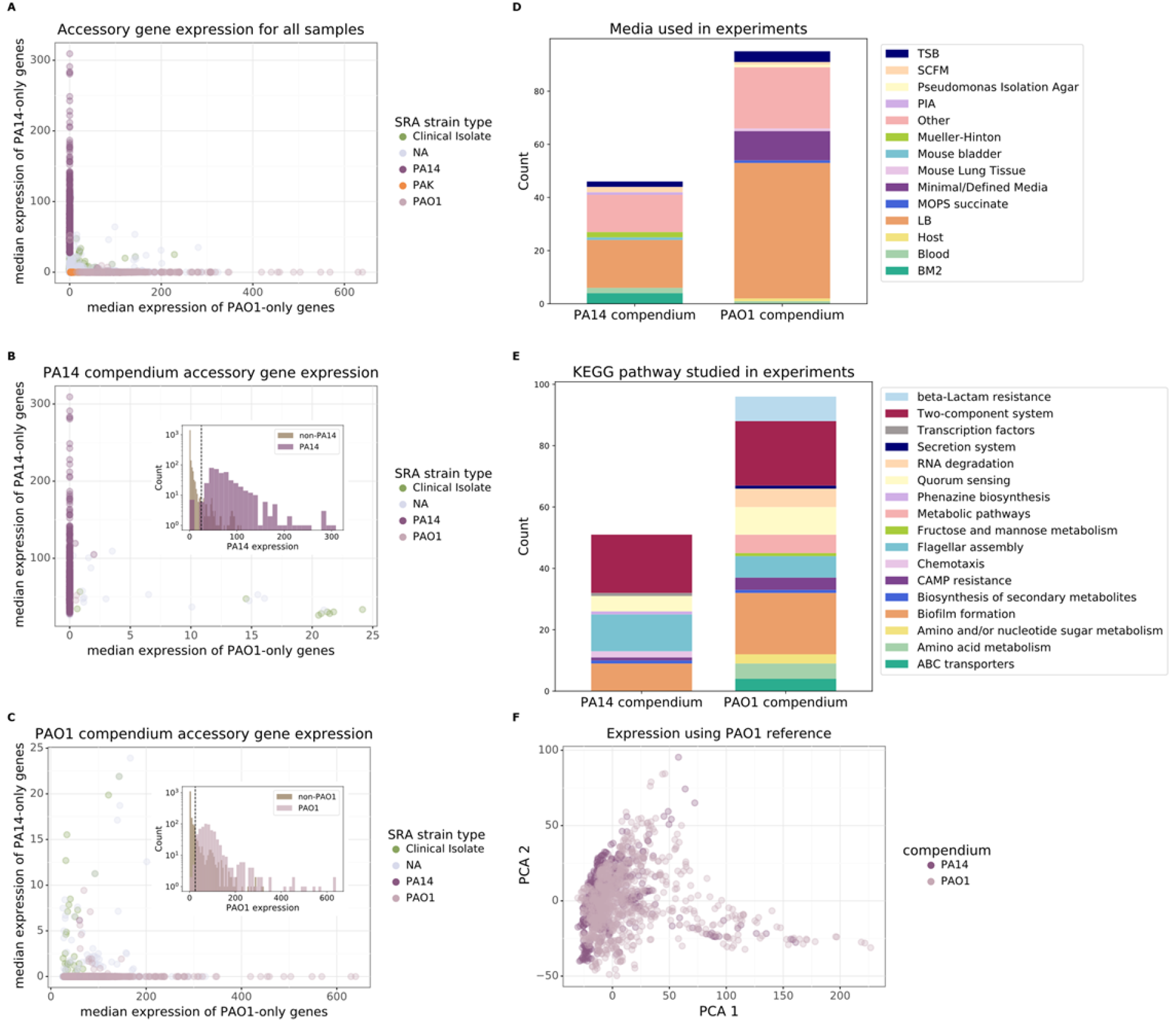
Creation and composition of PAO1 and PA14 compendia. A) Median accessory gene expression of all samples mapped to PAO1 and PA14 reference strains, as described in Doing *et al*., co-submitted manuscript B) Median accessory gene expression of samples in the PA14 compendium with the inset plot showing the distribution of accessory expression in PA14 SRA-annotated samples versus non-PA14 SRA-annotated samples. C) Median accessory gene expression of samples in the PAO1 compendium with the inset plot showing the distribution of accessory expression in PAO1 SRA-annotated samples versus non-PAO1 SRA-annotated samples. D) Media or groups of media used by experiments across the two compendia, E) The KEGG pathway(s) associated with genes perturbed (i.e., genes ‘knocked-out’ / ‘knocked-down’ or over-expressed) in experiments, compared across the two compendia. There are some experiments that did not perturb any gene and there are also some genes that did not have a KEGG annotation – these are not displayed. F) PCA of PAO1 and PA14 compendia aligned to PAO1 reference (the same structure is seen using the PA14 reference, data not shown).

Both the PAO1 and PA14 sample compendia represent diverse transcriptional phenotypes. Both compendia contained experiments using different media, with LB (51% of strain PAO1 experiments and 39% of strain PA14 experiments) as the predominant medium in both (Figure 1D). Many experiments relied on genetic manipulations that targeted genes related to multiple different functions and pathways (Figure 1E) but, interestingly, the KEGG pathway annotations for manipulated genes showed that the three largest categories for both the PAO1 and PA14 compendia were biofilm, quorum sensing and two component systems, suggesting commonalities among the functions of genes that researchers have interrogated in each strain.

Despite the genomic differences between PAO1 and PA14 strains, the differences in the transcriptome within each strain across many the contexts (e.g. growth conditions, genetic manipulations or other mutations) is much greater than the differences between strains (Figure 1F). The centroid difference between PAO1 transcriptomes and PA14 transcriptomes mapped to the PAO1 reference is 25 (30 using the PA14 reference, data not shown), which is smaller compared to the spread of the samples within each strain-specific compendium (6259 for PAO1 and 5126 for PA14 transcriptomes using the PAO1 reference for read mapping; 6125 for PAO1 and 6051 for PA14 transcriptomes using the PA14 reference for read mapping). This diversity among samples for each strain indicated the value of using these types of gene expression compendia to identify similarities and differences in transcriptional patterns between PA14 and PAO1 across the many contexts that have been studied.

### Certain core genes and pathways are transcriptionally stable across strain types

We first examined the transcriptional relationship between core genes. Since core genes are shared by both strain types, we asked: do core genes have similar correlation profiles between the two strain types? Because the PAO1 sample compendium and the PA14 sample compendium are comprised of different sets of experiments, we couldn’t directly correlate gene expression profiles so instead we assessed a second-order correlation to determine transcriptional stability across strains. Here we defined transcriptional stability as the similarity of transcriptional neighbors between homologous core genes in the two compendia -i.e. two homologous core genes are similar if they are most closely correlated with the same set of genes. To do so, we first linked PAO1 and PA14 homologs for all 5,349 core genes. For each core gene, we examined the correlation of their correlations to all other core genes across all samples in each of the strain-specific compendia. If two homologous core genes had a high second order correlation (also called *transcriptional stability)*, we considered those core genes to be highly stable (Figure 2B). After performing this pairwise comparison across all core genes, we defined a subset of genes that were *most stable* and another subset that were *least stable* by taking the top and bottom 5% of genes based on their transcriptional stability (Figure 2B, Table S1).

**Figure 2:**
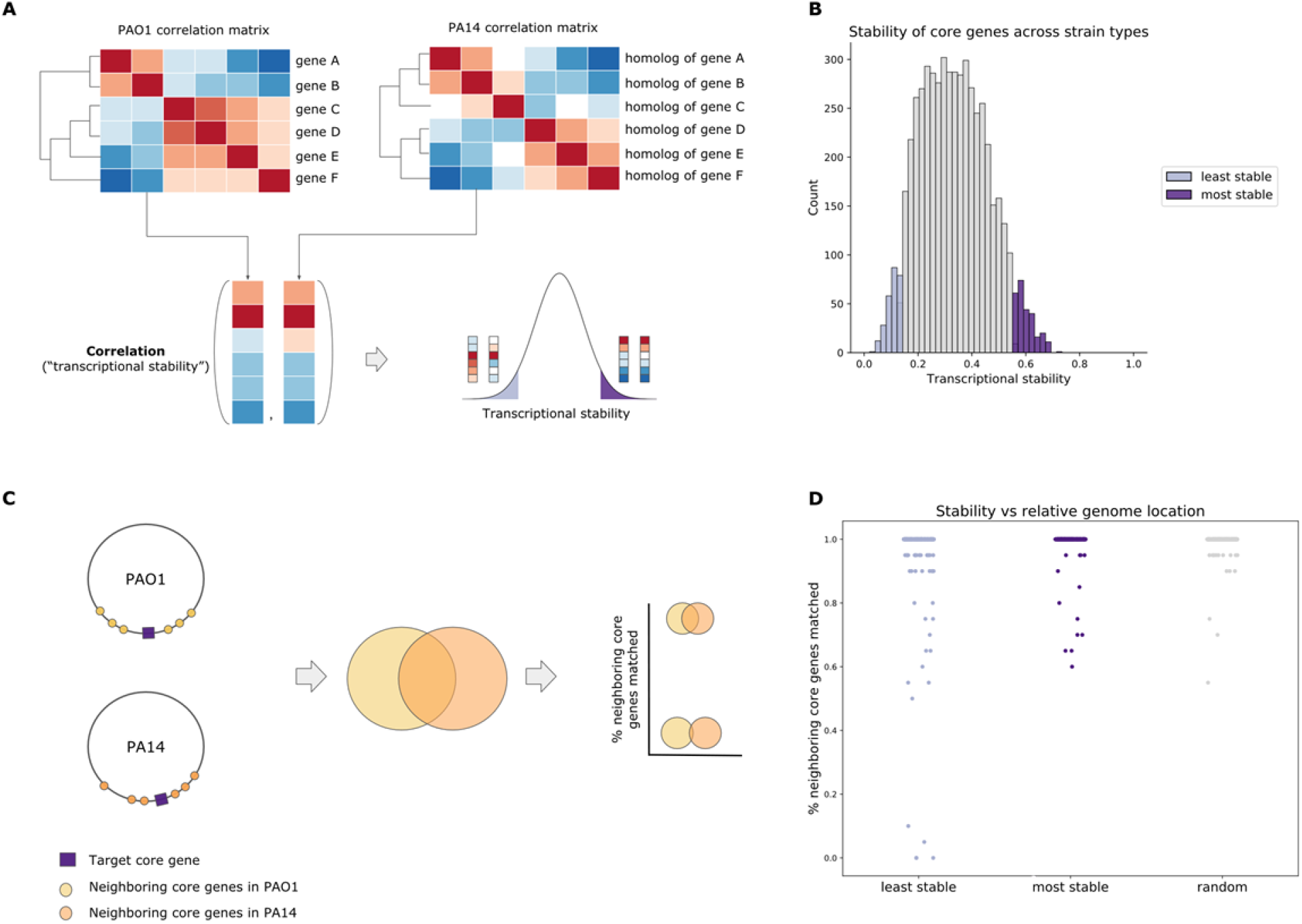
Relationships between core genes. A) Workflow describing how the most and least stable core genes were identified by comparing the correlation profile of homologous gene pairs. B) Distribution of correlation scores representing how stable core genes are across strains with most stable core genes highlighted in dark purple and least stable core genes in light purple. C) Workflow to test if least stable genes are located in different regions of the genome across strains. Given a target least stable or most stable core gene, we compare the overlap of neighbors in PAO1 versus the neighbors in PA14. If the overlap (% neighboring core genes matched) is high, meaning that many of the neighbors are the same in PAO1 and PA14, then this would indicate that the gene is located in the same neighborhood in both strains. D) Consistency of location of homologous genes in PAO1 versus PA14 strains in least stable core genes (light purple), most stable core genes (dark purple) and a random set of core genes (grey).

The most stable core genes were significantly enriched in association with pathways that represented essential functions: translation, RNA metabolism, DNA repair and recombination, central carbon metabolism (*sdhABCD, sucABCD, lpd*)^53^, amino acid metabolism, peptidylglycan biosynthesis (Table S2). These pathways were consistent with the functions of core essential genes identified by Poulsen *et al* (40% of most stable core genes were also found to be essential).^54^ In addition to these essential pathways, stable core genes also included genes within the type VI secretion system (T6SS)^55^ and genes in the Hcp Secretion Island-I (H1-T6SS). We confirmed that TVI genes were expressed in a significant fraction of conditions (Table S1-mean expression), demonstrating that these correlations likely represent true transcriptional stability between strains. However we were surprised to find T6SS genes amongst the most stable genes, given their diversity between strains. When we analyzed these genes more carefully, we found that there were numerous identical homologs within each genome which poses a problem for read mapping. Thus, these genes and other genes with identical paralogs were removed from further discussion (see Doing *et al*., co-submitted manuscript, Supplemental Dataset S1 for list of genes with identical or near identical paralogs).

The KEGG pathways significantly most enriched in least stable core genes were genes involved in the biosynthesis of paerucuramin (encoded by the *pvc* genes) which was interesting in light of the result that expression of the *pvc* gene regulator led to different product ratios in strain PA14 and strain PAO1 (Table S3).^56^ In addition, genes related to degradation of aromatic compounds including tyrosine and benzoate were also enriched in those genes less stable across strains (Table S3). Given the existence of least stable core genes, which are inconsistent in their co-expressed genes between strains PAO1 and PA14 (Figure 2B), this might suggest that this subset of core genes contributes to the difference in transcriptional regulation observed across lineages. For example, differences in virulence factor production was noted for PAO1 and PA14^57^ and this may be due to differences in the levels of aromatic amino acid catabolites that can influence the production of QS molecules and phenazines.^58,59^

Given that these least stable genes have inconsistent co-expressed neighbors across strain type, we hypothesized that one reason might be that these least stable genes are in different genomic neighborhoods and therefore are regulated differently between PAO1 versus PA14. To compare the neighborhood of homologous genes, we compared the percent overlap of neighboring core genes in PAO1 versus PA14. When we compared the neighborhood between homologous genes, this did not appear to explain differences in stability for most core genes, (Figure 2C, 2D). However, differences in genomic context could be a contributing factor for a subset of the least stable genes; genes located in different neighborhoods across strains include PA0982, PA2520 (*czcA*), PA3867, PA2226 (*qsrO*), and PA3507.^42^ Overall, this analysis suggests that a change in genome location was not the primary driver for instability in these least stable genes. Additionally, since co-operonic genes are co-transcribed, we might expect their “neighbors” (genes with whom they are most highly correlated) to be from the same operon, assuming their homologous genes are also organized into the same operon. Since transcriptional stability is determined based on neighbor congruency across strains, we checked that there wasn’t a bias such that stability could just be predicted based on whether a gene belonged to an operon and indeed even genes that are not in operons can still be stable and there isn’t a strong correlation between stability and operon size (Figure S1).

As a validation, we recalculated the most and least stable genes using the *P. aeruginosa* microarray compendium described in Tan et al^47^ (Figure S2A, Table S4). We found consistency between the transcriptional stability scores (R^2^=0.5, Figure S2B) and a significant (p-value=3.8e-48) over-representation of “most stable” core genes found using the array compendium within the most stable core genes found using the RNA-seq compendium. Again, the *pvc* genes were among the least stable (Table S4). It was also interesting to note that the gene-gene correlation scores within the microarray compendium were lower than for genes in the RNA-seq compendium.

**Figure S1:**
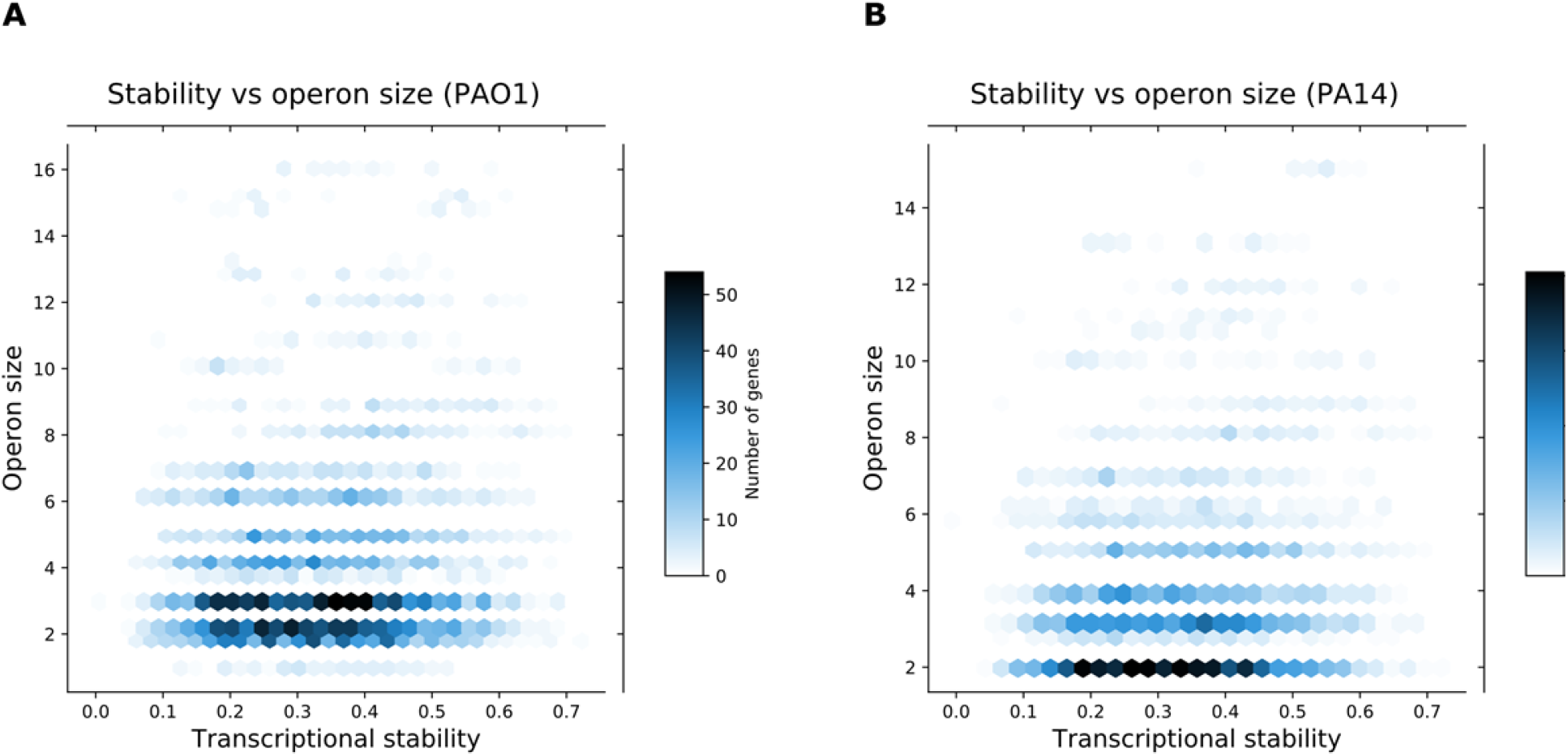
Effect of operon size on transcriptional stability. Correlation between transcriptional stability versus size of operons in A) PAO1 compendium and B) PA14 compendium.

**Figure S2:**
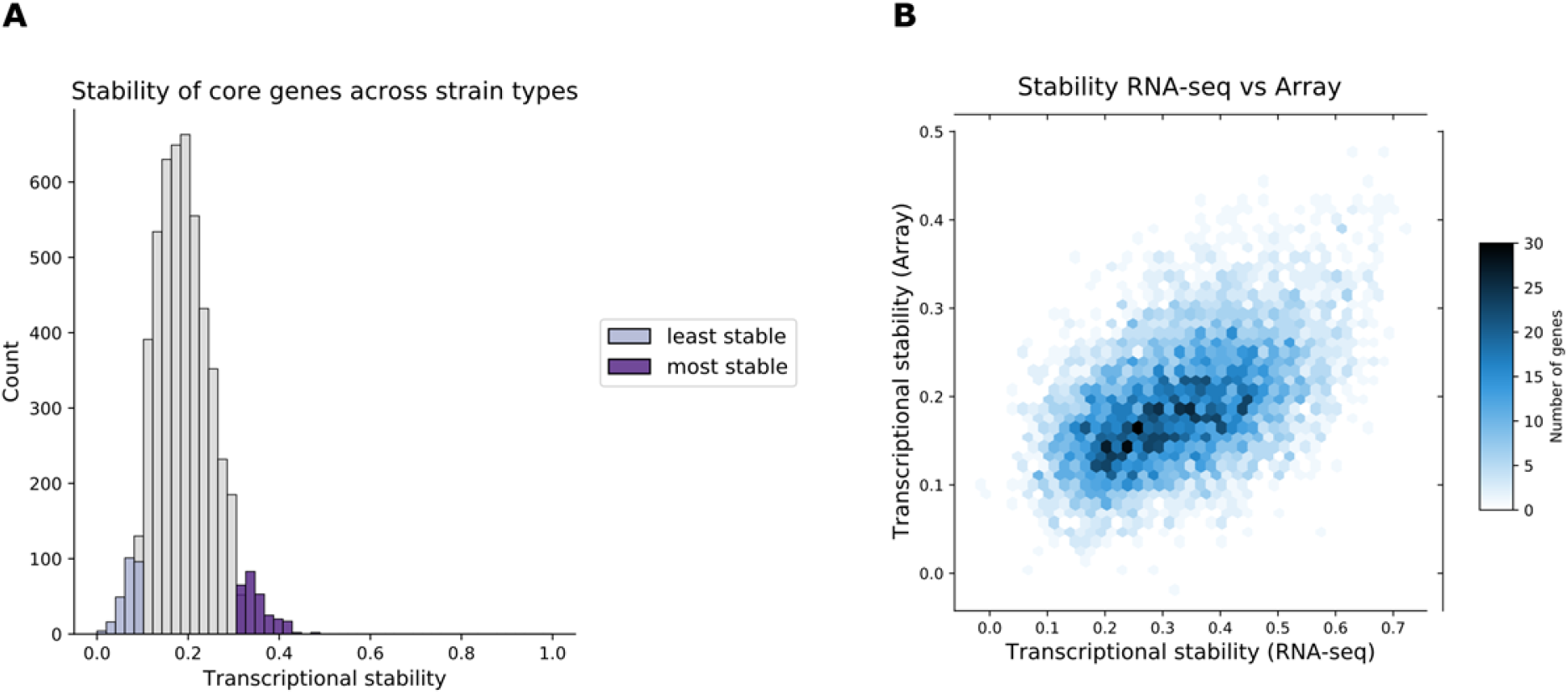
Validation of most stable core genes using an array compendium. A) Distribution of correlation scores representing how stable core genes are across strains with most stable core genes highlighted in dark purple and least stable core genes in light purple using the array compendium. B) Correlation of transcriptional stability values using the RNA-seq compendium versus the array compendium.

### Core genes that are less stable are more often co-expressed with accessory genes

It is suggested that accessory genes can be co-opted into existing core gene sets as well as influence their expression, which can lead to niche adaptations.^60–62^ Therefore, we hypothesized that accessory genes might alter the transcriptional stability of core gene expression, and help to explain the difference between the most and least stable core genes. We tested this hypothesis by determining whether global transcriptional stability was associated with co-expression with accessory genes. For each core gene in the most and least stable sets, we identified the 10 most co-expressed genes in each strain. We then calculated the proportion of these co-expressed genes that were accessory genes compared to the proportion of accessory genes in the genome (Figure 3A). We found that most stable core genes tended to exhibit less co-expression with accessory genes – most stable core genes have fewer transcriptional neighbors that are accessory genes (Figure 3B).

**Figure 3:**
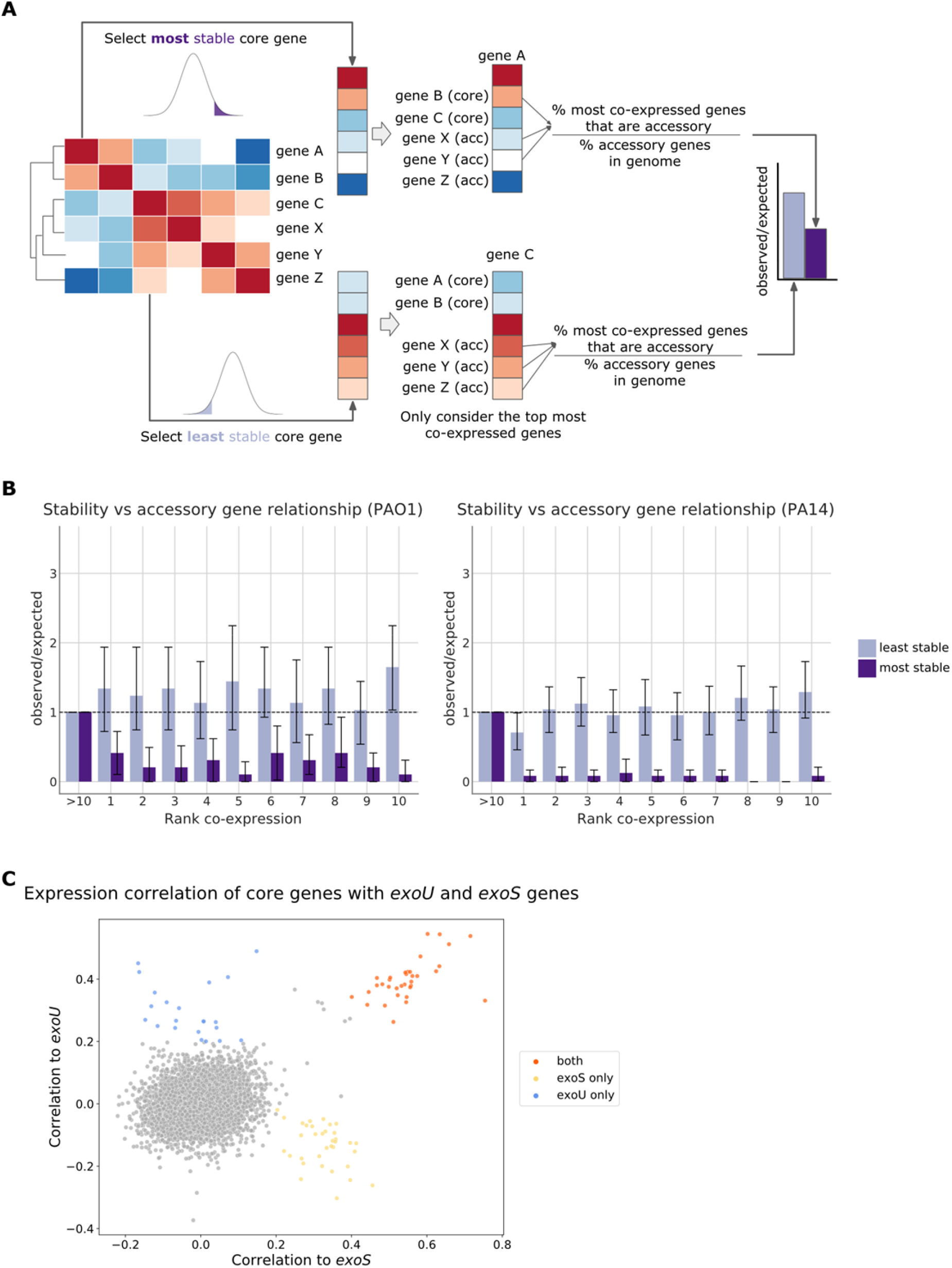
Relationships between core and accessory genes. A) Workflow describing how genes co-expressed with most and least stable core genes were identified. Given a most stable core gene, the topmost co-expressed genes were identified and the proportion of co-expressed genes that are accessory compared to the expected proportion of accessory genes in the genome was calculated. B) The proportion of accessory genes that the most stable (dark purple) and least stable (light purple) core genes were co-expressed with. C) Scatterplot depicting the correlation between core genes and two T3SS accessory genes, *exoS* versus *exoU*.

### Comparison of exoU and exoS co-regulated genes

We also examined transcriptional stability in the context of the type III secretion system (T3SS), which is a virulence mechanism that allows *P. aeruginosa* to inject toxic effector proteins into the cytoplasm of eukaryotic cells. We selected the T3SS because, while there are four effector proteins secreted by this system (ExoU, ExoS, ExoY and ExoT), where the gene encoding ExoU and ExoS are strain-specific (i.e. the presence of *exoU* and *exoS* tends to be mutually exclusive in PAO1 and PA14 strains), and they have differential effects on the host.^37,63^ We found 34 core genes that were highly co-expressed with both *exoU* and *exoS*. Some of these were related to the T3SS secretion machinery (*popNBD, pcrGDVH, pscCN)* as well as genes encoding the effector protein *exoT* (Figure 3C). In addition to core genes that were highly co-expressed with both *exoU* and *exoS*, we found some core genes that were highly expressed with only *exoU* or only *exoS*: 22 core genes were highly co-expressed with *exoU*, while 35 core genes were highly co-expressed with *exoS*. Core genes more highly co-expressed with *exoS* were related to metabolic pathways, including *acoRB* which is involved in acetoin catabolism, and *adh*, an alcohol dehydrogenase (Table S5), and *spcS*, which encodes an ExoS chaperone. In contrast, genes that were more highly co-expressed with *exoU* included *oprD*^64^, which encodes an outermembrane porin implicated in carbapenem resistance, a trait associated with ExoU positive strains.^65^ Another gene more co-expressed with *exoU* is *rhlR*, which encodes a regulator involved in quorum sensing in response to a signal generated by the co-expressed synthase RhlI, and *pheC*, which encodes a periplasmic cyclohexadienyl dehydratase involved in phenylalanine biosynthesis. Interestingly, *pheC* is adjacent to the *rhlR-rhlI* genes. Despite ExoS and ExoU being secreted by the same secretion pathway, we identified distinct groups of core genes that might suggest different environments in the two clades are found –acetoin, a major product of the Enterobacteriaciae, may be present when theT3SS is activated for many ExoS-containing strains. These different functional modes are consistent with the genes having different pathogenic effects on the host cell function.^38^ Overall, most stable core genes tend to be less co-expressed with accessory genes, suggesting that the interrelationship between core and accessory gene expression depends on strain type. Therefore, including both the core and accessory genes will be important for revealing possible new regulatory relationships in different strain types.

### Clustering by co-expression patterns identified accessory-accessory modules

We used a graph-based clustering approach, called Affinity propagation^66^, to cluster accessory genes into modules based on their correlation profiles. In the PAO1 compendium, the 202 accessory genes were clustered into 28 modules (Supplementary Table S6), with a median size of 5 genes. Similarly, the 530 PA14 accessory genes were clustered into 70 modules, with a median size of 7 genes (Supplementary Table S7). Based on a manual inspection, we found that accessory modules tended to correlate with known operons as expected; the predicted operon for each gene is listed and manual inspection highlights that this method correctly identified the co-regulation of co-operonic genes. This finding was expected as we confirmed that co-operonic genes were highly correlated in expression with each other in these compendia (Doing *et al*., co-submitted manuscript) and in most cases operon structure is conserved across strains. Indeed, while some accessory modules contained co-operonic genes (e.g. module 13 (*wbp* genes involved in O-antigen biosynthesis) and module 23 (*psl* genes involved in extracellular matrix production) Table S6), most modules contain genes from more than one operon. There were multiple modules that contained an uncharacterized or partially characterized transcriptional regulators with non-co-operonic genes which will provide the basis for future mechanistic exploration such as modules 10, 19 (which contains *vqsM*), 22, and 24 in strain PAO1 (Table S6) and modules 6, 24 and 28 in strain PA14 (Table S7). We also observed that some accessory modules had a similar function in PAO1 and PA14, such as those related to fimbrial biosynthesis and LPS biosynthesis. It is worth noting that module 45 for strain PA14 which was enriched in genes that may be problematically mapped due to the presence of high identity paralogs (see Doing *et al*., co-submitted manuscript) and caution should be used in considering this module as it may reflect a technical feature of the data rather than a biological one.

We have provided these tables of accessory gene modules as a resource for scientists to explore and guide future research. For example, there are many missing gene annotations, particularly for accessory genes.^67^ Given that the most recent CAFA (Critical Assessment of Functional Annotation) study found that co-expression data was highly predictive of function, these modules could point to possible candidate functions that can be experimentally verified.^68^ In general, these supplementary tables (Table S6, S7) can provide a summary of the landscape for what accessory genes have been researched and what remains to be explored. The expression statistics provided could also be used to help scientists to filter the set of genes to focus on. Another possible application of these tables is to facilitate operon discovery. In general, there exist very few microbes with experimentally verified operons and these experiments are expensive and time-consuming to perform at a genome-wide scale. Thus computational operon prediction methods are important.^69^ While more validation is necessary, the results of our co-expression clustering approach found many modules that captured co-operonic genes which might suggest a possible contribution to existing prediction methods.

## Discussion

Through our analysis of core and accessory gene expression, we discovered that there is diverse transcriptional behavior within the different gene groups between strains of *Pseudomonas aeruginosa*. Despite core genes being shared across strains, core genes differ in their transcriptional patterns with some being highly stable and others less so. The most stable core genes are less co-expressed with other accessory genes. One possibility is that accessory genes might change the transcriptional regulation of these core genes which is why most stable genes are less co-expressed with accessory genes.^70^ For example, it is known that an insertion sequence, which is a type of accessory gene, can influence the expression of an existing gene via the existing promoter or by the formation of a hybrid promoter.^60–62^ Additionally, a study found that accessory genes could modify the function of core genes between two strains of *Streptococcus pneumoniae*.^70^ Overall, by using the compendia of expression data, we were able to distinguish types of core genes by their global transcriptional patterns that will help inform future studies of regulatory mechanisms to better understand *P. aeruginosa* strain diversity.

In this analysis we focused on examining core and accessory genes within *P. aeruginosa*, however, this analysis can be extended to other gene groups or organisms beyond *P. aeruginosa*. For example, bacteriophage genes, which are viral genes that are integrated into the bacterial genome, are another type of gene contained within the *P. aeruginosa* genome. Similar to the other gene groups, changes in phage expression have been shown to affect phenotypes. For example, upregulation of phage genes was found to promote biofilm development.^71^ Investigating phage gene expression within the transcriptional landscape would require generating a suitable reference genome that includes all phage genes, which doesn’t currently exist and may be challenging to generate given the rapid rate of phage sequence evolution.^72^ This work could also be extended to other two-strain type models in different microbes to help understand the phenotypic heterogeneity. Comparisons between the two lineages of *Staphylococcus aureus* in Australia with different resistance phenotypes^73^ and the two *Plasmodium vivax* lineages with different geographical origins^74^ may be well suited to this model. Other examples include comparing two groups of strains based on phenotypes: comparing rare versus common *Escherichia coli* strain types^75,76^, comparing *Staphylococcus aureus* resistant versus sensitive strains^77^, comparing *Acinetobacter baumannii* outbreak in different hospital wards^78^, or comparing symbiotic and nonsymbiotic species^79^.

One limitation is that our analysis is limited to examining two strain types at a time. However, there exist additional clades that we might want to consider. For example, in *P. aeruginosa*, there exist many clinical strains in addition to PAO1 and PA14. To apply our approach to include clinical strains, we would need to generate a clinical reference genome to get a more accurate expression of clinical accessory genes. We also need to determine the core and accessory annotations in order to account for an additional strain type. Finally, we need to determine a new metric to assess stability since we will have more than two variables for our correlation calculation.

By leveraging expression compendia with hundreds to thousands of samples, this study reveals the complex relationship among and between core and accessory genes that should be considered when designing future regulatory analyses to further study *P. aeruginosa* diversity.

## Methods

### PAO1 and PA14 expression compendia

For these analyses we used the PAO1-mapped and PA14-mapped RNA-seq compendia described in Doing *et al*., co-submitted manuscript^42^ Doing *et al*. provided a filtered and median-ratio (MR) normalized compendia containing 2,333 samples mapped to 5,563 genes using cDNA sequences from the PAO1 reference genome (PRJNA331)^80–82^ and 5,891 genes using cDNA sequences from the PA14 reference genome (PRJNA386)^21^. The filtering steps included removal of sparse samples (i.e. samples having a high number of genes with zero counts) since having many undetected genes indicated some technical issue. They also discarded samples based on too high or low median expression of housekeeping genes, which are genes that we expect to be consistently expressed at relatively high levels across samples. Median-ratio (MR) normalization was then performed to enable comparisons across samples. The expression data can be found in the following repository: https://osf.io/vz42h/.

Although SRA provides annotations for strain type, we used the expression activity of the accessory genes (i.e. genes that are specific to PAO1 or specific to PA14) to bin samples into PAO1 and PA14 compendia. There were several advantages to using the expression activity instead of the metadata provided by SRA. First, this approach allowed us to create a pipeline that can easily extend to new datasets where metadata isn’t available. Despite the metadata being available in this case, the strain names require curation. Next, we were able to leverage more data as opposed to losing ∼10 -22% of the samples which have an unknown strain annotation. Last, by using the expression activity we can obtain a cleaner separation where we avoid samples with high levels of both PAO1 and PA14 accessory expression. A sample is considered PAO1 if the median gene expression of PA14 accessory genes is less than 25 MR normalized estimated counts and PAO1 accessory genes is greater than 25 MR normalized estimated counts. Similarly, a sample is considered PA14 if the median gene expression of PA14 accessory genes is greater than 25 MR normalized estimated counts and PAO1 accessory genes is less than 25 MR normalized estimated counts. A threshold of 25 is used based on the distribution of accessory genes in SRA-annotated PAO1 samples compared with the distribution of SRA-annotated non-PAO1 samples. We defined the threshold to be one that separated between these two distributions. After applying these thresholds, we obtained a PAO1 sample compendium containing 890 samples and 5,563 genes. Similarly, the PA14 sample compendium contained 505 samples and 5,892 genes. These two compendia contained experiments that used a variety of different media and interrogated different genes. This associated metadata was curated using the same process as described in Doing *et al*., co-submitted manuscript, where the GEOquery R package was used to collect the metadata associated with the studies contained in the compendia.

### Core and accessory annotations

By definition, core genes are those genes that are present in all strains. While accessory genes are those that are present in at least one strain. Here, we used the annotations of core genes obtained from the BACTOME website^52^, where core genes are those that had at least 90% sequence homology between the two strain types and accessory genes are those remaining genes that are either PAO1 or PA14 specific. There are 5,361 core genes and 202 PAO1 and 530 PA14 accessory genes respectively.

### Distribution of PAO1 and PA14 sample compendia using principal components analysis (PCA)

Samples from the PAO1 sample compendium (890 samples) and PA14 sample compendium (505 samples) were selected (1,395 samples) from the original filtered and normalized RNA-seq compendium (2,333 samples) that was mapped to both the PAO1 and PA14 reference. Our selected resulted in two datasets with 1,395 samples aligned to the PAO1 and PA14 references. We used the sklearn library to compress the selected PAO1 and PA14 datasets into the first 2 principal components. We then calculated the centroid of the PAO1 and PA14 sample compendia by taking the average of the first 200 principal components since 90% of the variance was explained by 200 components. Then we measured the pairwise Euclidean distance between the two centroids to get the difference in mean between two strain distributions. To calculate the spread of the PAO1 and PA14 sample compendia we summed the variance for the first 200 principal components.

### Correlation matrix

When we generated a Pearson correlation matrix using the MR normalized counts, we found that many gene pairs had a high correlation because many genes are related to the same pathway and therefore have a similar expression profile. This is consistent with Myers *et al*.^83^, who found that there can be an over-representation of genes associated with the same pathway (i.e. a large fraction of gene pairs represent ribosomal relationships). This very prominent signal makes it difficult to detect other signals. To remove this very dominant global signal in the data, we first log10-transformed the expression data and then applied the signal balancing technique called SPELL^84^. The SPELL algorithm calculated the SVD of the expression matrix to get the factorized set of matrices. We then applied the Pearson correlation on the SPELL processed matrix, which is the factorized gene coefficient matrix *U*. This coefficient matrix represents how genes contribute to independent latent variables that capture the signal in the data where the variance of the variables is 1. The correlation of the SPELL matrix relates genes based on the gene coefficient matrix. In other words, the correlation of the SPELL matrix relates genes on their contribution to singular vectors (SV) which capture linear relationships between genes. A high correlation means that a pair of genes contributes similarly to a singular vector, which are the axes pointing in the direction of the spread of the data and capture how genes are related to each other. The advantage of using SPELL is that the gene contributions are more balanced so that redundant signals (i.e., many genes from the same pathway or genes that vary together) are represented by a few singular vectors as opposed to many samples. More balanced also means that more subtle signals can be amplified (i.e., genes related by a smaller pathway are also captured by a similar number of SVs as larger pathways). However, the one caveat is that SPELL can amplify noise - i.e., an SV that corresponds to some technical source of variability now has a similar weight to other real signals.

### Transcriptional stability of core genes

For this analysis we started with a PAO1 core gene, say PA0001. First, we selected the correlation scores for all genes related to PA0001. Next, we selected the homologous PA14 gene, PA14_00010, and likewise pulled the correlation scores for how genes were related to PA14_00010. Then, we calculated the Pearson correlation between the two correlation profiles to obtain a *transcriptional stability* score. We repeat this analysis for all core genes. We removed 5 PAO1 core genes and 10 PA14 core genes that had ambiguous mapping across strains (i.e., they mapped to multiple gene ids). High scores (genes in the top 5%) indicated that the transcriptional relationships of the core genes were consistent or *most stable*. Whereas low scores (genes in the bottom 5%) indicated the core genes were *least stable*. This analysis was performed using both the RNA-seq compendium (this study) and the array compendium described in Tan *et al*.^47^ The processing steps described above were applied to both compendia.

### Core gene stability versus location consistency

For each most stable core gene, we selected the neighboring core genes (10 genes upstream from the most stable core gene and 10 genes downstream) in PAO1 and PA14. We then calculated the overlap between the 20 neighboring core genes in PAO1 and PA14 and asked: how many of the 20 neighboring core genes in PAO1 are homologous to the neighboring genes in PA14? If there was a large overlap between the neighboring genes across strains, that would indicate that the most stable core gene was located in the same genomic region in PAO1 and PA14. We repeated this calculation for all most stable core genes. We also performed this calculation starting with the least stable core genes.

### KEGG enrichment analysis

The goal of an enrichment analysis is to detect coordinated changes in prespecified sets of related genes (i.e., those genes in the same pathway or share the same GO term). Here we used the one-sided Fisher’s exact test. This test asked if there a significant over-representation of KEGG pathways in the most and least stable core gene sets. Specifically, this method tested if the proportion of KEGG annotations within the most stable core gene set was more than expected compared to the proportion of KEGG genes in the total dataset. The pathways used in this analysis were downloaded from the KEGG website (http://www.genome.ad.jp/kegg/) using the python Bio.KEGG library. These pathways can be found in the associated repository:

https://github.com/greenelab/core-accessory-interactome/blob/master/3_core_core_analysis/pao1_kegg_annot.tsv and https://github.com/greenelab/core-accessory-interactome/blob/master/3_core_core_analysis/pa14_kegg_annot.tsv

### Relationships between core and accessory genes

Here we examined which genes the most and least stable core genes are related to. For this analysis, we start with the most stable core genes and ask: is the highest correlated gene core or accessory? For a given stable core gene, we got a list of genes sorted by their correlation score, making sure to remove any genes that were co-operonic with the start gene. The operon data was provided by Geoff Winsor and includes computationally predicted annotations from DOOR^85^ as well as curated annotations from PseudoCAP^81,82^. We repeated this selection and filtering for all most stable core genes so that for all most stable core genes we have a list of the top 10 most co-expressed genes. Then we can calculate the proportion of the most co-expressed genes that are accessory. This proportion is normalized by the proportion of accessory genes within the whole genome. The resulting score is near 1.0 if the proportion of accessory gene relationships is no different compared to the baseline proportion of accessory genes in the genome. If the resulting score is higher or lower than 1.0 then the proportion of accessory gene relationships are more or less than expected compared to baseline. We repeated this calculation for the second most co-expressed genes, the third most co-expressed genes, and so forth so that we have a fold change over random value for each top 10 co-expressed ranking. We also performed this same calculation for the eleventh most co-expressed genes and beyond to act as a baseline. This analysis is repeated starting with least stable core genes as well.

### Accessory-accessory module detection

To get accessory-accessory modules we started with the gene expression of only accessory genes, which included 202 PAO1-specific genes for the PAO1 sample compendium and 530 PA14-specific genes for the PA14 sample compendium. To define a set of modules we first applied Pearson correlation on the SPELL-processed expression matrices (see description above) to get correlation scores for how similar each pair of genes are based on their transcriptional profiles. Then we applied clustering on the correlation matrices. The clustering method used is called affinity propogation^66^. Affinity propagation is a graph-based clustering algorithm similar to k Means, but without the need to specify the number of clusters a priori. Affinity propagation finds “exemplars” which are members of the input (or genes in our case) that are representative of clusters. These exemplars are equivalent to “centroids” in k-means. Exemplars are determined by an exchange of messages, which consists of “responsibility” and “availability”. Responsibility quantifies how well suited a gene X is to be an exemplar based on how similar other genes are to X and how available X is. Availability measures how likely a gene is to choose X as its exemplar. Availability is calculated as the responsibility of X with itself and the responsibility of X towards all other genes. The clustering returned 28 modules for PAO1 and 70 modules for PA14. We validated that those accessory modules tended to correlate with known operons based on a manual inspection. While we did not perform a thorough evaluation to determine that this clustering approach is the best, we found that the modules appear to be biologically relevant and can serve as a steppingstone for future research.

### Software

All scripts used in these analyses are available in the GitHub repository (https://github.com/greenelab/core-accessory-interactome) under an open-source license to facilitate reproducibility of these findings (BSD 3-Clause). The repository’s structure is described in the Readme file. The virtual environment was managed using conda (version 4.9.2), and the required libraries and packages are defined in the environment.yml file.

## Acknowledgements

The authors would like to thank Geoff Winsor for providing operon data that we used for our analysis relating core gene to other accessory genes. The authors would also like to thank Jake Crawford and Natalie Davidson for reviewing the software associated with this work and providing valuable feedback. This work was supported by grants from the Gordon and Betty Moore Foundation (GBMF4552 to CSG), Cystic Fibrosis Foundation (HOGAN19GO to DAH, GREENE21GO to CSG and STANTO19R0 to support SLN) and the Flatley Foundation. Finally, to the National Institutes of Health (NIH) through NIDDK P30-DK117469, P30 DK117469 and R01 HL151385.

## Author Contributions

AJL: Formal analysis; Investigation; Methodology; Project administration; Software; Visualization; Writing-original; Writing-review

GD: Data curation; Writing-review

SLN: Data curation; Writing-review

TR: Writing-review

DAH: Conceptualization; Funding acquisition; Methodology; Supervision; Writing-original, Writing-review

CSG: Conceptualization; Funding acquisition; Methodology; Supervision; Writing-review

## Supplementary Tables

**Table S1:** Transcriptional stability of all core genes.

**Table S2:** Most stable core genes and their associated KEGG pathways.

**Table S3:** Least stable core genes and their associated KEGG pathways.

**Table S4:** Transcriptional stability of core genes using PAO1 RNA-seq compendium versus PAO1 array compendium.

**Table S5:** Core genes correlated with *exoS* and *exoU*.

**Table S6:** Membership of all PAO1 accessory genes to modules.

**Table S7:** Membership of all PA14 accessory genes to modules.

